# Heterospecific song quality as social information for settlement decisions: an experimental approach in a wild bird

**DOI:** 10.1101/748103

**Authors:** Jennifer Morinay, Jukka T. Forsman, Blandine Doligez

## Abstract

Assessing local habitat quality via social cues provided by con- or heterospecific individuals sharing the same needs is a widespread strategy of social information use for breeding habitat selection. However, gathering information about putative competitors may involve agonistic costs. The use of social cues reflecting local habitat quality acquired from a distance, such as acoustic cues, could therefore be favoured. Bird songs are conspicuous signals commonly assumed to reliably reflect producer quality, and thereby local site quality. Birds of different species have been shown to be attracted to breeding sites by heterospecific songs, but whether they can use fine heterospecific song features as information on producer (and by extension habitat) quality remains unknown. We used a playback experiment in a wild population of collared flycatchers (*Ficedula albicollis*), a species known to eavesdrop on dominant great tits’ (*Parus major*) presence and performance, to test whether flycatchers preferred to settle near broadcasts of a high quality great tit song (i.e. song with large repertoire size, long strophes, high song rate), a low quality great tit song or a chaffinch song (control). Among old females, aggressive ones preferred to settle near broadcasts of high quality tit song and avoided broadcasts of low quality tit song, while less aggressive females preferred to settle near broadcasts of low quality tit song. Male personality or age did not influence settlement decisions. Our results show that collared flycatcher females use great tit song quality features as information for settlement decisions, though differently depending on their own competitive ability and/or previous experience with great tit songs. Our study therefore further illustrates the complex condition-dependent use of heterospecific social information for breeding habitat selection.

## Introduction

When habitat quality varies in time and space, choosing where to breed can have crucial consequences for individual fitness. Hence, strong selective pressures can be expected to promote behavioural strategies allowing individuals to optimize habitat selection decisions. In particular, individuals can collect and use information about habitat quality to choose among alternative breeding sites or patches (Danchin et al. 2004, Dall et al. 2005). Such information can be acquired from the individual’s own interactions with its environment, i.e. its personal experience (‘personal information’, e.g. its own reproductive success, Switzer 1997). Alternatively, information can be acquired from observing the interactions of other individuals sharing similar needs (either con- or heterospecific putative competitors) with the environment and the result of these interactions, either inadvertently or when they intentionally communicate with others (“social information”, Danchin et al. 2004, Dall et al. 2005).

When cueing on others, individuals can rely on the mere presence of con- or heterospecifics (patch / site occupancy or density, Thiebault et al. 2014), an information that can be easily accessible but does not directly inform on the fitness consequences of others’ decisions. Individuals can also use the performance of others, i.e. the success obtained after making a decision, an information that can be more difficult to access but better informs about the fitness consequences of the decision. In the context of breeding habitat selection, information about others’ performance (when available) can often be used only after a delay, up to a whole breeding season (Boulinier et al. 2008). When breeding synchrony with individuals sharing similar needs is low, eavesdropping on the reproductive investment of early competitors could inform on habitat quality for decisions later in the same season (e.g. Seppänen et al. 2011, Forsman and Seppänen 2011, Loukola et al. 2013). However, assessing competitors’ performance can involve proximity to their breeding sites and therefore increase the risk of agonistic interactions (e.g. Merilä and Wiggins 1995, Forsman et al. 2018). Therefore, in this case, balancing the trade-off between information accuracy and reliability, on the one hand, and information availability and costs associated to information gathering, on the other hand, may require reducing such costs. This should favour the use of cues reflecting others’ performance obtained to a low cost, such as cues obtained from a distance.

Among such cues, acoustic signals have been shown to be an information source easily eavesdropped on, even from a long distance (e.g. anti-predatory strategies involving eavesdropping on conspecific and heterospecific alarm calls; reviewed in Magrath et al. 2015). Experimental studies have clearly shown that calls and songs can, on their own, induce conspecific (Hahn and Silverman 2006) and heterospecific attraction (Fletcher 2008, Szymkowiak et al. 2017) to otherwise empty breeding sites, a property sometimes used in reintroduction programmes to enhance local settlement of released animals (e.g. Ward and Schlossberg 2004). Importantly, signals used in sexual communication, which include acoustic signals, are selected (1) to be conspicuous, allowing emitters to be detected by the highest possible number of potential partners (in intersexual communication) and/or competitors (in intrasexual communication), and (2) to reliably reflect individual quality (e.g. in terms of health, competitive ability, etc.; Andersson 1994, Catchpole and Slater 2008). Female birds have for example been shown to eavesdrop on males singing contests and adjust mate choice and reproductive behaviour accordingly (Otter et al. 1999, Mennill et al. 2002). Therefore, acoustic signals could provide social information on individual quality (Møller 1991, e.g. Buchanan and Catchpole 2000, Bischoff et al. 2009), and thereby indirectly on habitat / territory quality. Such acoustic information use could occur not only within, but also between species, which has not been explored yet.

Using a playback experiment in a wild population of collared flycatchers *Ficedula albicollis*, we experimentally tested whether individuals use songs from heterospecific competitors as a source of information for nest site selection and whether they modulate the use of this cue depending on song features, reflecting the quality of its emitter. Migratory flycatchers are known to use different heterospecific social information from their main competitor, the resident great tit *Parus major*, for nest site selection (tit presence: Kivelä et al. 2014; tit density: Forsman et al. 2008; tit early reproductive investment: Seppänen et al. 2011, Loukola et al. 2013; tit breeding phenology: Samplonius and Both 2017). Prospecting tit nests to gather information on tit presence or reproductive investment may nevertheless be risky (Merilä and Wiggins 1995, Forsman and Thomson 2008, Forsman et al. 2018). Therefore, flycatchers could be expected to rely also on less costly cues, such as great tit songs, which can be heard from a distance and whose characteristics (repertoire size and strophe length) have been shown to correlate with great tit quality (McGregor et al. 1981, Lambrechts and Dhondt 1986).

Upon flycatchers’ arrival from migration, we broadcasted artificially created great tit songs of either high quality (large repertoire, long strophes, high song rate) or low quality (small repertoire, short strophes, lower song rate) and monitored flycatchers’ settlement in experimental zones. If flycatchers are attracted by great tits songs when choosing where to breed, they should settle preferentially in zones with broadcasted tit songs; in addition, if flycatchers use information about great tit quality as reflected by song features, they should settle preferentially in zones with broadcasted high quality tit songs, i.e. presumably indicating high quality habitat. We also tested whether the choice of a zone with high vs. low quality tit song depended on flycatchers’ age, which may affect previous experience with great tit songs, and aggressiveness, which may affect the ability to face competitive costs with great tits. Finally, we tested whether flycatchers adjusted early reproductive investment, as previously found in this population in response to tit density (Forsman et al. 2008), here according to the experimental song treatment.

## Materials and methods

### Study area and population monitoring

The experiment was conducted in spring 2017, in a patchy population of collared flycatchers breeding on the island of Gotland (Sweden, Baltic Sea). In this population, collared flycatchers and dominant great and blue tits (*Cyanestes caeruleus*) partly share the same ecological niche: they are hole-nesting species breeding in tree cavities and readily accept nest boxes provided in excess in the study area (Doligez et al. 2004c). Collared flycatchers start arriving from migration on the breeding grounds late April-early May, i.e. 2 weeks on average after the beginning of tit breeding season. Every year, approx. 1/3 of the nest boxes available in the population is occupied by collared flycatchers, 1/3 by tits (among which 3/4 are great tits) and 1/3 remains empty. In all nest boxes occupied by flycatchers, we captured females during incubation and males during the chick rearing period (for nests reaching this stage). All captured individuals were identified (or ringed if previously unringed), measured, weighed and aged based on plumage criteria (yearlings vs. older adults; Svensson 1992). Nest boxes were then visited regularly throughout the breeding season to record the main breeding variables for each breeding pair (laying and hatching date, clutch size, number and condition of nestlings, final fledging success).

### Playback experimental design

In the 13 forest patches chosen for the study on the basis of a sufficiently high number of nest boxes (> 30, and up to 180), we established experimental zones composed of five neighbouring nest boxes (exceptionally four in one zone), with four boxes (exceptionally three boxes in one zone) spread around a central nest box (at approximately 20 m distance); experimental zones were separated from each other by at least 40 m (i.e. each zone was surrounded by at least one row of non-experimental nest boxes). Each selected forest patch contained 3 to 9 experimental zones, for a total of 57 experimental zones (19 of each broadcast treatment). We conducted the playback experiment between the 29^th^ of April and the 27^th^ of May, i.e. during the whole period of flycatcher settlement. During these 29 days, we broadcasted at the centre of each experimental zone either (i) a great tit song track with high quality song features, i.e. mimicking singing activity of high quality individuals (i.e. large repertoire, long strophes, high song rate, McGregor et al. 1981, Lambrechts and Dhondt 1986, Rivera-Gutierrez et al. 2010), (ii) a great tit song track with song features of low quality individuals (i.e. with a small repertoire size, short strophes, low song rate) or (iii) a song track from a species with no a priori influence on flycatcher settlement decisions, i.e. not sharing the same ecological niche: the chaffinch, *Fringilla coelebs*, as a control. The proximity of experimental zones reliably reflected the natural density of great tit breeding pairs in our forest patches. Furthermore, experimental zones of the three treatments were balanced within each forest patch as far as possible. Because tit and chaffinch songs can be heard from a long distance (more than 100 m in our forest patches), a prospecting flycatcher should thus have been able to simultaneously hear several experimental broadcasts and choose among treatments.

Song tracks were broadcasted from dawn (1 hour before sunrise) for a duration of 17 hours, corresponding to dusk at the beginning of the broadcasting period and up to 1h30 before dusk at the end of the broadcasting period. Along the broadcasting period, the starting hour of the broadcast was adjusted (15 minutes earlier every 10 days) to match the seasonal change of dawn hour, but the length of the track remained unchanged. To match the natural singing activity of great tits, we broadcasted 10 minute-long song periods every 30 minutes from dawn to 3 hours after dawn, and then every hour till the end of the sequence (see Figure A1), similarly to Krebs et al. (1978). Tracks were broadcasted at ~85-95dB, i.e. close to the natural sound amplitude of great tit songs (McGregor and Horn 1992; sound amplitude checked at 1 m distance with a sound level meter “Dr. Meter MS10”). In each experimental zone, song track was broadcasted from a camouflaged loudspeaker (Zealot S1) attached 1.5-2 m above ground on a tree next to the central nest box of each experimental zone.

### Playback song structure

To create the broadcasted sound tracks while avoiding pseudoreplication, we used songs from 4 different great tits to mimic songs of high quality tits, from 4 others to mimic songs of low quality tits and from 4 different chaffinches for controls. Great tit songs were recorded in the same population in 2016, at dawn chorus, with a Sennheiser MKH70 microphone and a Zoom H4N recorder. Chaffinch songs were recorded either on Gotland in 2016 (1 individual) or on the Swedish mainland (3 individuals) and available on-line (Xeno Canto on-line database, www.xeno-canto.org, accessed in April 2017; recordings ID: XC84011, XC196974, and XC27602). Each sound track was composed of songs originating from only one individual to mimic the presence of a single singing individual in each experimental zone, and in each zone, the song track broadcasted remained unchanged during the whole broadcasting period, to avoid mixing signals from different individuals in case flycatchers were able to individually recognize singers. All recordings were in .wav format to ensure sufficient sound quality and had a sample frequency of 44.1 kHz and a resolution of 16 bit. Using Audacity software (v. 2.1.0, http://audacity.sourceforge.net/), original recordings were high-pass filtered with a threshold below the song minimum frequency (2 kHz), modified to create the song bouts (see Figure A1) and amplified. We amplified whole song bouts (see Figure A1) but kept natural variations in amplitude within bouts, to mimic singing bird movements to a flycatcher listening from a fixed point.

### Controlling for neighbouring live great tits

To keep nest boxes in the experimental zones available to flycatchers and avoid songs from live great tits to strongly interfere with our broadcasted great tit songs, we prevented great tits (but not blue tits) to settle in our experimental zones from early April by narrowing nest box entrance hole of each box to 28 mm diameter. At the beginning of the broadcasting period, i.e. on the 29^th^ of April, we expanded again the nest box entrance hole to 32 mm diameter (recommended size for both great tits and flycatchers, L. Gustafsson pers. comm.). As a consequence, late blue and great tits could also settle in the experimental zones during the broadcasting period. When this happened before the first flycatcher pair had settled in the experimental zone and the nest box density was sufficiently low (for 3 of the 57 zones), we slightly relocated the experimental zone by adding one nest box on the edge of the zone and excluding the box occupied by the tit pair, in order to provide the same number of available nest boxes (5) to the first flycatchers to settle in all zones.

The singing performance of neighbouring great tits might locally affect our treatment. Therefore, we controlled for the singing activity of tits within or in close vicinity to each experimental zone. We counted the number of different songs that could be heard close-by from the tree where the loudspeaker was placed, during 10 minutes picked at random before 10 am and in between two broadcasted songs. We recorded this measure of singing activity by alive great tits for each zone during 4 to 7 days a week depending on the experimental zone and field time constraints, obtaining between 16 and 27 measures per zone, and we averaged it over the whole broadcasting period to obtain a measure of mean song ‘bias’ in each zone.

### Aggressiveness test

We estimated the aggressiveness level of flycatchers settling in the experimental zones during nest building or early egg laying stages. We followed the protocol detailed in Morinay et al (2019). In short, at the beginning of the test, an observer attached (i) clay decoys representing either a flycatcher pair or a male great tit to the nest box of the focal pair and (ii) a loudspeaker broadcasting songs of the corresponding species below the nest box. The observer then sat camouflaged 8-10 m away and recorded all the behaviours of both the male and the female flycatchers for 15 minutes if both individuals were seen at least 5 minutes during this first period, or up to 25 minutes if at least one of them arrived at the end of the first 15 min only, to allow describing flycatchers’ behaviour for at least 5 minutes. We conducted one test with flycatcher decoys and one with great tit decoy. However, if one individual was not seen during either test, we conducted more tests (up to 5), with a day break between two consecutive tests. To avoid pseudoreplication, we used 10 sets of flycatcher decoys, 10 sets of great tit decoys, 5 different song tracks per species, and we randomized the song track used with a given decoy set. Aggressiveness score was then later estimated as the number of moves within 2 m away from the nest box (between branches or to the box, as well as attacks on decoys) plus the number of chases performed against live intruders, standardized per 15 minutes (a measure whose repeatability within and between years is around 0.25, Morinay et al. 2019). Over the 99 flycatcher pairs that started building nests in our experimental zones, we obtained aggressiveness data for 92 females and 95 males, among which 78 females and 61 males were later captured and aged, for 84 nests with eggs in total.

### Statistical analyses

We first tested whether overall flycatcher settlement in experimental zones differed between song broadcast treatments (high quality great tit song, low quality great tit song, chaffinch song) by comparing nest box occupancy probability between treatments with a 3-sample test for equality of proportions, using the ‘prop.test’ function in R.

Second, among settled pairs, we tested whether the probability for flycatchers to settle in a given song broadcast treatment depended on individual and environmental factors using multinomial mixed models. We fitted generalized linear models with the broadcast treatment of the zone chosen by each flycatcher (3 modalities) as the response variable and included as fixed effects individual’s age and aggressiveness score, settlement date, the presence of previously settled great tits and flycatchers (as shown by the presence of nest material in a box; 2 separate binary variables) in the experimental zone on the day of choice and mean song bias. We included age and settlement date, as well as their two-way interaction, because late arriving birds and yearlings have been found to rely more on social information from great tits compared to early arriving and older ones (Seppänen and Forsman 2007). We included the presence of settled great tits and flycatchers prior to settlement of the focal bird, as well as mean song bias, to control for social attraction. We also included the two-way interaction between aggressiveness and age, as well as the interactions between either aggressiveness or age and the presence of flycatchers, to account for potential age- and personality-specific responses in the use of social information, as previously shown in this species (Morinay 2018). Because of our limited sample size and to avoid over-parametrization, we could not include all interactions at once and thus tested them one-by-one. We initially included the forest patch as a random effect, but removed it as the variance associated to this effect did not differ from zero (based on the Heidelberger and Welch’s half-width test, ‘heidel.diag’ R function). We could not include playback track as a random effect because of model convergence issues.

Finally, we tested whether flycatchers adjusted early reproductive investment, measured here by (i) laying date, (ii) the delay between settlement and laying, and (iii) clutch size, according to the broadcast treatment (3 modalities) using (generalized) linear mixed effects models. Besides the treatment, we included as random effects the forest patch and as fixed factors the same effects as mentioned above, except for settlement date, which was included only in the model for clutch size.

### Linear model implementation

Even though nest box choice and reproductive investment are likely to be joint decisions by both pair members, we fitted separate models for males and females, because combining the effects of both sexes in a single model leads to a reduced sample size (more females are captured than males).

We implemented Bayesian linear models with the *MCMCglmm* function (“MCMCglmm” R package, Hadfield 2010). We centred and scaled all continuous explanatory variables to ease comparison. Besides the two-way interaction terms, which were removed when the 95% Credible Interval encompassed zero, we did not select models for fixed effects (Mundry and Nunn 2009). To account for a potential bias in the capture of individuals with respect to personality types (Garamszegi et al. 2009), we fitted the same models including individuals whose aggressiveness score was estimated but that were not captured later on, i.e. models without the age effect; the results remained qualitatively unchanged (results not shown). We implemented models fitting the treatment chosen using the ‘categorical’ family (two parametrizations depending on the interaction term included: 15 or 35 × 10^5^ iterations, burn-in =1 or 3 × 10^4^, thinning-interval = 700 or 1700); we fixed the variance-covariance residual matrix to 2/3 for the diagonal terms (variance) and 1/3 for all the off-diagonal terms (covariance; Hadfield 2016). We implemented the models for laying date with the Gaussian family (12 × 10^4^ iterations, burn-in = 6 000, thinning interval = 50) and for the delay in laying and clutch size with the Poisson family (11 or 15 × 10^5^ iterations, burn-in = 1 or 2 × 10^4^, thinning interval = 500 or 700 depending on the interaction term included). For all models, we used normally distributed priors with a mean of 0 and a large variance (10^8^) for fixed effects and inverse-Gamma priors for the residual (when not fixed) and random variances. Model convergence was assessed visually and by comparing three chains per model using the *gelman.diag* and *gelman.plot* functions (Gelman & Rubin approach, Plummer et al. 2006). Autocorrelation levels were always below 0.1 and effective sample sizes above 1500 per chain.

### Animal welfare note

To minimize disturbance during the aggressiveness tests, we approached the nest box as quietly as possible and hided below a camouflage net. The Ringing Centre from the Museum of Natural History in Stockholm granted permission for catching and ringing adults (here 77 females and 60 males) with individually numbered aluminium rings (licences nb. 471:M025 to JM and 471:M043 to Cécile Vansteenberghe). We captured male and female adults in the nest, either directly (females during incubation) or using swinging-door traps (both parents during chick rearing). We set up traps for 30-60 minutes maximum depending on nestling age (30 min when nestlings were 5 days old or younger), to avoid nestling starvation if parents did not resume feeding during the catching period; we checked upon the traps every 5-10 minutes, and removed as soon as the adults had been caught. We started catching sessions after 6:00 am to let birds feed and provision nestlings undisturbed for at least two hours after the night period (sunrise occurs approx. at 4 am during spring). We handled adults during 5 to 10 minutes and released them straight after manipulation or (when catching both parents during nestling feeding) kept them until capturing the partner (up to 40 min maximum). All manipulations were done in accordance with the Swedish legislation applying at the time.

## Results

### Occupancy pattern according to broadcast treatment

The probability for a nest box to be chosen by collared flycatchers did not differ between treatments: the total number of settled pairs was 33 in the high quality great tit song treatment, 27 in the low quality tit song treatment and 39 in the chaffinch song (control) treatment (χ^2^_2_ = 3.55, p-value = 0.17).

### Individual and environmental effects on the choice of a broadcast treatment

The probability to choose an experimental zone of a given broadcast treatment depended on female aggressiveness score but differently for yearling and older females (interaction aggressiveness score by age in Table 1). Among older females, less aggressive ones were more likely to settle in zones of the low quality great tit song treatment (relation between aggressiveness level and probability to settle in the low quality song vs. control treatment: estimate [95% CI]: −3.47 [−6.45; −0.96]; Table 1, Figure 1B), while more aggressive ones were more likely to settle in zones of the high quality great tit song treatment (relation between aggressiveness level and probability to settle in the high quality song vs. control treatment: estimate [95% CI]: 1.33 [0.14; 2.63]; Table 1, Figure 1A). The probability to choose an experimental zone of a given broadcast treatment also depended on the presence of competitors previously settled in the zone: flycatchers were more likely to settle in a great tit song treatment zone (either high or low quality compared to control) when other great tits had previously settled in the same zone (Table 1); they were also more likely to settle in a control zone than in a high quality great tit song zone when other flycatchers had previously settled in the experimental zone (Table 1). Neither settlement date, song bias, male age, nor male aggressiveness affected the probability to settle in a zone of a given broadcast treatment (Table 1).

**Table 1.**
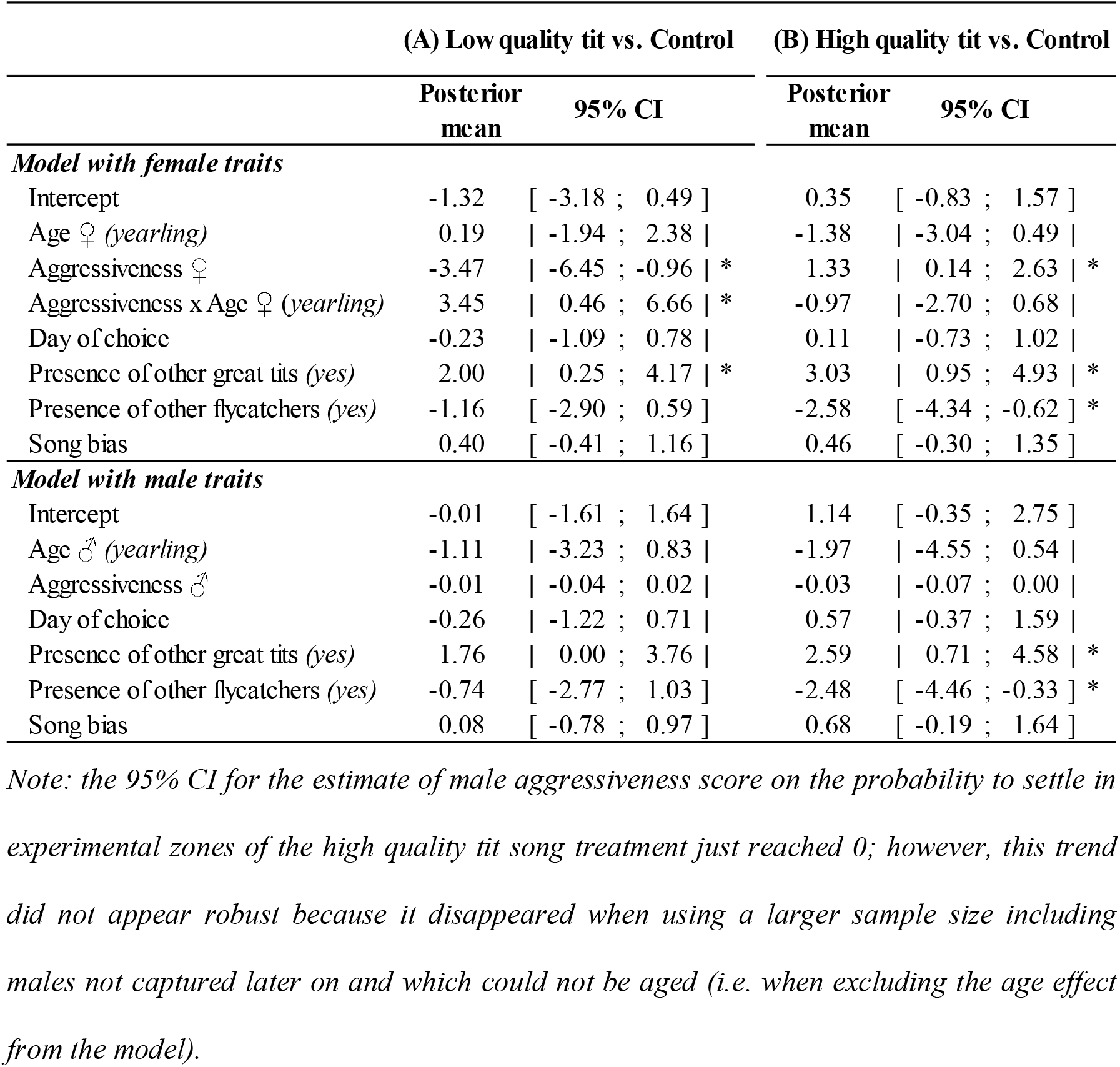
Effect of individual and environmental variables on the probability to settle in experimental zones of a given broadcast treatment. Model outputs are estimates (posterior means and 95% Credible Intervals) for settlement in an experimental zone of (A) the low quality great tit song vs. the control (chaffinch song) treatment and (B) the high quality great tit song vs. the control treatment (i.e. the control treatment served as reference here), for females and males separately (see text). For qualitative traits, the estimated category is given in parentheses. Stars indicate estimates whose 95% CI do not overlap zero.

|  | (A) Low quality tit vs. Control |  | (B) High quality tit vs. Control |  |
| --- | --- | --- | --- | --- |
|  | Posterior mean | 95% CI | Posterior mean | 95% CI |
| ***Model with female traits*** |  |  |  |  |
| Intercept | -1.32 | [ -3.18 ; 0.49 ] | 0.35 | [ -0.83 ; 1.57 ] |
| Age ♀ (*yearling*) | 0.19 | [ -1.94 ; 2.38 ] | -1.38 | [ -3.04 ; 0.49 ] |
| Aggressiveness ♀ | -3.47 | [ -6.45 ; -0.96 ] * | 1.33 | [ 0.14 ; 2.63 ] * |
| Aggressiveness x Age ♀ (*yearling*) | 3.45 | [ 0.46 ; 6.66 ] * | -0.97 | [ -2.70 ; 0.68 ] |
| Day of choice | -0.23 | [ -1.09 ; 0.78 ] | 0.11 | [ -0.73 ; 1.02 ] |
| Presence of other great tits (*yes*) | 2.00 | [ 0.25 ; 4.17 ] * | 3.03 | [ 0.95 ; 4.93 ] * |
| Presence of other flycatchers (*yes*) | -1.16 | [ -2.90 ; 0.59 ] | -2.58 | [ -4.34 ; -0.62 ] * |
| Song bias | 0.40 | [ -0.41 ; 1.16 ] | 0.46 | [ -0.30 ; 1.35 ] |
| ***Model with male traits*** |  |  |  |  |
| Intercept | -0.01 | [ -1.61 ; 1.64 ] | 1.14 | [ -0.35 ; 2.75 ] |
| Age ♂ (*yearling*) | -1.11 | [ -3.23 ; 0.83 ] | -1.97 | [ -4.55 ; 0.54 ] |
| Aggressiveness ♂ | -0.01 | [ -0.04 ; 0.02 ] | -0.03 | [ -0.07 ; 0.00 ] |
| Day of choice | -0.26 | [ -1.22 ; 0.71 ] | 0.57 | [ -0.37 ; 1.59 ] |
| Presence of other great tits (*yes*) | 1.76 | [ 0.00 ; 3.76 ] | 2.59 | [ 0.71 ; 4.58 ] * |
| Presence of other flycatchers (*yes*) | -0.74 | [ -2.77 ; 1.03 ] | -2.48 | [ -4.46 ; -0.33 ] * |
| Song bias | 0.08 | [ -0.78 ; 0.97 ] | 0.68 | [ -0.19 ; 1.64 ] |

*Note: the 95% CI for the estimate of male aggressiveness score on the probability to settle in experimental zones of the high quality tit song treatment just reached 0; however, this trend did not appear robust because it disappeared when using a larger sample size including males not captured later on and which could not be aged (i.e. when excluding the age effect from the model).*

**Figure 1.**
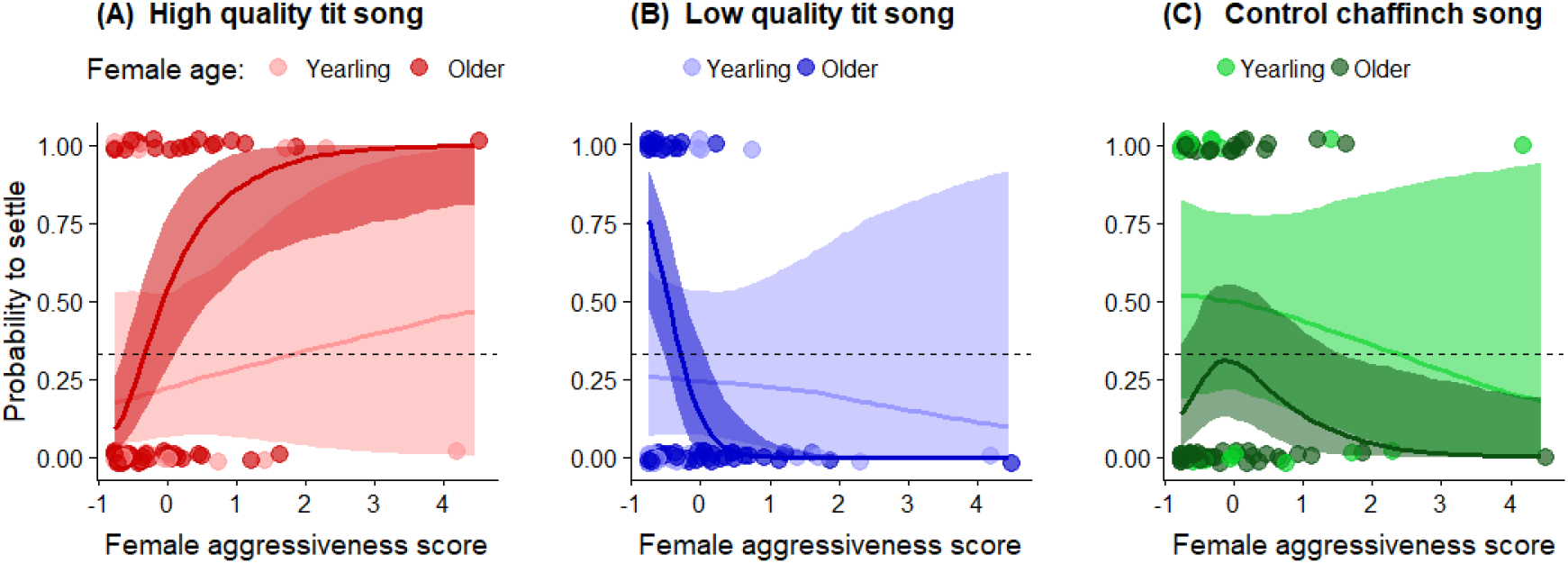
**Probability for flycatchers to settle in an experimental zone according to the broadcast treatment (3 modalities, A, B, C) depending on female aggressiveness score and age** (light red/blue/green: yearling females; dark red/blue/green: older females). Predicted mean probabilities (solid lines) and their 95% Credible Intervals (shaded areas) were derived for average estimates of all other model parameters. Points indicate actual settlements (1 for settlement in the broadcast treatment considered, 0 for settlement in one of the other two treatments). The horizontal dashed line represents the probability to settle at random in a zone of the broadcast treatment considered (i.e. 0.33).

### Early reproductive investment according to broadcast treatment

We found no difference between broadcast treatments in flycatcher laying date, delay between settlement and laying, and clutch size (all 95% CI encompassed zero, Table A1). As expected, yearling females were found to lay eggs later in the season than older females (Table A1). Aggressiveness scores and male age had no effect on early reproductive investment (Table A1; all 95% CI for the two-way interactions encompassed zero, results not shown).

## Discussion

Flycatcher settlement in experimental zones depended on aggressiveness level in interaction with female age: among old females, more aggressive ones settled preferentially in zones with high quality tit songs, while less aggressive preferred zones with low quality tit songs. This result provide the first evidence for eavesdropping on heterospecific song features related to emitter’s quality in addition to song presence itself. This source of social information did however not affect flycatchers’ early reproductive investment (laying date, clutch size), suggesting that different information sources are used for different breeding decisions (Doligez et al. 2008), and calling for a finer understanding of the fitness benefits of using each information source.

### Why and when using great tit song features for settlement decisions?

Migratory flycatchers may not have easy access to information about breeding habitat quality when returning from migration and have been shown to rely on resident, already settled, great tit presence and early reproductive investment for their own settlement decisions under strong time constraints (e.g. Forsman and Seppänen 2011, Kivelä et al. 2014). Songs may allow flycatchers to easily assess great tit density, but also quality, from a distance, while direct information about early reproductive investment might be more difficult and costly to gather (Merilä and Wiggins 1995, Forsman et al. 2018, Samplonius and Both 2019).

Song production is overall costly (in terms of time, energy, predation risk and agonistic contests) and should thus be selected to honestly inform on producer’s quality (Gil and Gahr 2002), for example reflecting its past (Bischoff et al. 2009) or present parasitic load (Møller 1991, Buchanan and Catchpole 2000). Song features in great tit males have been shown to inform on male survival and reproductive success (McGregor et al. 1981, Lambrechts and Dhondt 1986, Rivera-Gutierrez et al. 2010), on mate quality (to females) during escalating song contests (Otter et al. 1999), and on competitive ability (to rivals) at the conspecific level (Peake et al. 2005). Great tit song features have not directly been related to territory quality (Lambrechts and Dhondt 1988, but see Hoi-Leitner et al. 1995, and Manica et al. 2014 for other species), but great tit males singing longer strophes were found to be dominant at feeders (Lambrechts and Dhondt 1986) and more willing to engage in territorial defence (McGregor and Horn 1992). Overall, these studies suggest that great tit song features revealing singers’ high quality are likely to be associated with the acquisition and defence of a high quality territory, but also a better exploitation of habitat during nestling provisioning and nest defence against predators via increased vigilance and risk-taking (Krams 1998). Thus, cueing on great tit song features may be a relatively cost-free proximate mechanism for flycatchers to identify and select high quality individuals close to which it can be beneficial to settle (Forsman et al. 2002). Our experimental results confirmed the direct use of this cue by flycatchers for small-scale settlement decisions because we used broadcasted songs alone, i.e. decoupled from the actual presence of settled great tit pairs. This did not preclude flycatchers to simultaneously use the presence of great tit pairs in addition to the broadcasted songs, as reflected by higher settlement probability in presence of previously settled great tit pairs.

Nevertheless, the availability of great tit songs to newly arrived flycatchers may vary both within and between years. When flycatchers arrive on the breeding grounds, a large part of great tit females can have initiated incubation and thus great tit male singing activity can be largely reduced (Mace 1987, Amrhein et al. 2008). The time delay between great tit settlement and flycatcher arrival, as well as the time interval between the arrival of the first and last flycatchers, may strongly constrain the possibility for flycatchers to eavesdrop on great tit song and may emphasize the use of other information sources. The timing of great tit reproduction but also the synchrony of flycatcher arrival show high variability between years in this and other populations (Morinay et al. 2018, Samplonius and Both 2019), affecting the availability of cues linked to great tit presence and reproductive activity upon flycatcher arrival. Thus selective pressures should favour flexibility in the use of the different heterospecific cues in response to environmental variation. Great tit song characteristics may be used for flycatcher settlement decisions when tits are late and by early arriving flycatchers, while other information about tit quality and reproductive investment (e.g. clutch size, tit incubating or provisioning activity; Seppänen et al. 2011, Samplonius and Both 2017) or conspecific social information should be favoured otherwise, even though settlement date had no influence here on broadcast treatment choice. Manipulating information availability through the timing of song broadcast would be needed to explore this hypothesis. In our study, tit laying date was intermediate (34.8 ± 7.8 (SD), counted from the 1st of April) compared to other years (26.6 ± 6.9 for 2016, an early year, and 42.3 ± 3.9 for 2013, a late year; see Morinay et al. 2018), which may explain relatively small differences between broadcast treatments.

### Role of female, but not male, experience and competitive ability

Old females were able to use the information about both the presence and quality of great tit individuals provided by songs to select a breeding site. Yearling females were not impacted by song quality. This difference between yearling and older females may result from different past experience with great tit songs. Cueing on great tit song features may indeed require experience obtained only during the first breeding season for flycatchers, because just fledged flycatchers are usually not exposed to great tit songs before leaving on migration. The age difference in the use of a source of social information use could also be related to the lower competitive ability of yearling individuals, as already suggested for males in this population (Doligez et al. 1999). Flycatcher females may adjust their settlement decisions depending on the balance between benefits in terms of habitat quality and costs in terms of competition level: old and more aggressive females settled in the apparently most favourable habitats (as reflected by higher quality great tit songs) where they could cope with potentially higher competition level; old but less aggressive females still preferred settling near great tits, i.e. in habitats of supposedly higher quality than control (chaffinch song) zones, but they avoided zones where competition with great tits was expected to be highest. This could be in line with previous intraspecific results showing higher settlement of great tits near broadcasts of great tit songs with smaller repertoires, i.e. reflecting potentially lower quality individuals (Krebs et al. 1978): later-settlings individuals could indeed be low competitive individuals more prone to avoid potential competitive costs.

Alternatively, flycatcher females may have adjusted their response to our aggressiveness test after settlement depending on the apparent competitive level of neighbouring great tits. Indeed, we measured aggressiveness during nest building, at a time when playback songs were still broadcasted for most flycatcher pairs, and higher singing performance was suggested to induce social aggression, at least at the intraspecific level (Gil and Gahr 2002). Thus old females settled near apparently high quality great tits may have shown a higher aggressive response to intruders in response to such increased heterospecific social stimulation. In our population, aggressiveness score shows relatively high variability within individuals allowing for behavioural adjustment depending in particular on the context (Morinay et al. 2019). Nevertheless, both mechanisms, i.e. an adjustment of settlement choice according to aggressiveness or an adjustment of aggressiveness response according to songs of surrounding tits, involve female ability to discriminate low and high quality great tit songs and to adjust behaviour accordingly.

Interestingly, male age and aggressiveness score did not influence pair settlement with respect to the broadcast treatment. Even though nest site selection is a joint behaviour by both pair members, this could suggest that only female flycatchers were capable of adjusting their behaviour in response to great tit songs, which could reflect a higher ability to discriminate fine song features compared to males. Selective pressures may be higher in flycatcher females compared to males for the use of songs in the context of species recognition (hybridization with sympatric pied flycatchers in our population; Veen et al. 2001) and/or mate selection (higher constraints on females than males due to facultative polygyny; Gustafsson 1989). Former studies have shown differential auditory processes between sexes (Williams 1985, Negro et al. 2000), upon which selection could act differently. Alternatively, males could discriminate fine song features just as well as females (as suggested by widespread ‘dear enemy’ effects; Moser-Purdy and Mennill 2016), but may be less prone to use this information for settlement decisions if other social cues are more relevant at the spatial scale of male choice, supposedly involving smaller scales compared to females (Greenwood 1980, Doligez et al. 2004b, Arlt and Pärt 2008, Samplonius and Both 2017b, Morinay et al. 2018). Further work would be needed to assess whether flycatcher males can discriminate great tit song features and in this case which other information sources would be more valuable to them compared to females.

Overall, our results shed further light on the complexity of social information use by providing evidence for the use of refined heterospecific information sources such as the quality-related information contained in heterospecific acoustic signals for settlement decisions, with potential implications for songbird community dynamics. Further work is needed to assess how and when different information sources are used for different breeding decisions (see e.g. Doligez et al. 2008) depending on individual and environmental conditions, including the quantification of fitness benefits of using each information source in a given context.

## Acknowledgements

We are grateful to Lars Gustafsson for advice, to the land owners of Gotland for letting us work on their properties, and to the students and assistants who helped implementing this experiment (and its pilot in 2016), especially Cécile Vansteenberghe. We also thank Clémentine Vignal, Thierry Lengagne and Franck Perret for experimental and technical advices. JM was funded by the Ministère de l’Enseignement Supérieur et de la Recherche and by the Department of Ecology and Genetics from Uppsala University (PhD grant); JM was also funded by research grants from Uppsala University (Stiftelsen för Zoologisk Forskning, and the Liljewalsh foundation), the Animal Behaviour Society, the French Embassy in Sweden (ÖMSE program) and by the University of Lyon (IDEX mobility grant). JTF was funded by the Kone Foundation, and BD by the Centre National pour la Recherche Scientifique.

## Authors contribution

JM, JF and BD designed the experiment. JM collected and analysed the data, and drafted the manuscript. JM, JF and BD critically revised the manuscript. JM is the guarantor of this work and, as such, had full access to all of the data in the study and takes responsibility for the integrity of the data and the accuracy of the data analysis.

## Ethics

Permission for catching and ringing adult and young birds was granted by the Ringing Centre from the Museum of Natural History in Stockholm (licences nb. 471:M025 to JM and 471:M043 to Cécile Vansteenberghe). Aggressiveness tests only required observing individuals from a distance from below a camouflage net without catching them to avoid detrimental effect of aggressiveness test on the reproductive success of flycatchers (see the Methods section).

---

**Table A1.**
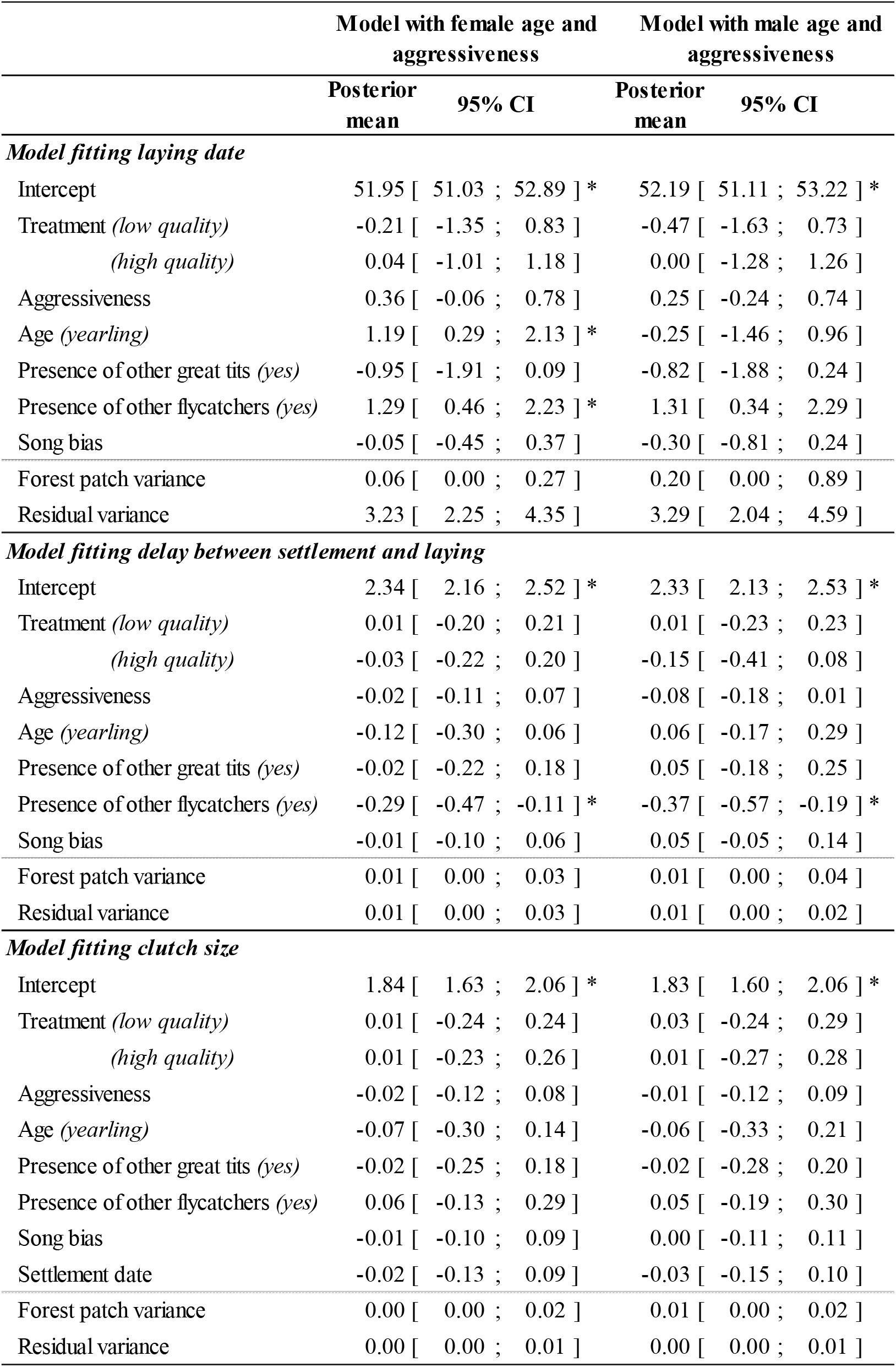
Output of models fitting flycatchers’ early-reproductive investment including either female or male age and aggressiveness score. Estimated levels are given in parentheses; older females, the control treatment, and the absence of other great tits or flycatchers serve as references. Stars indicate estimates which 95% CI do not overlap zero.

**Figure A1.**
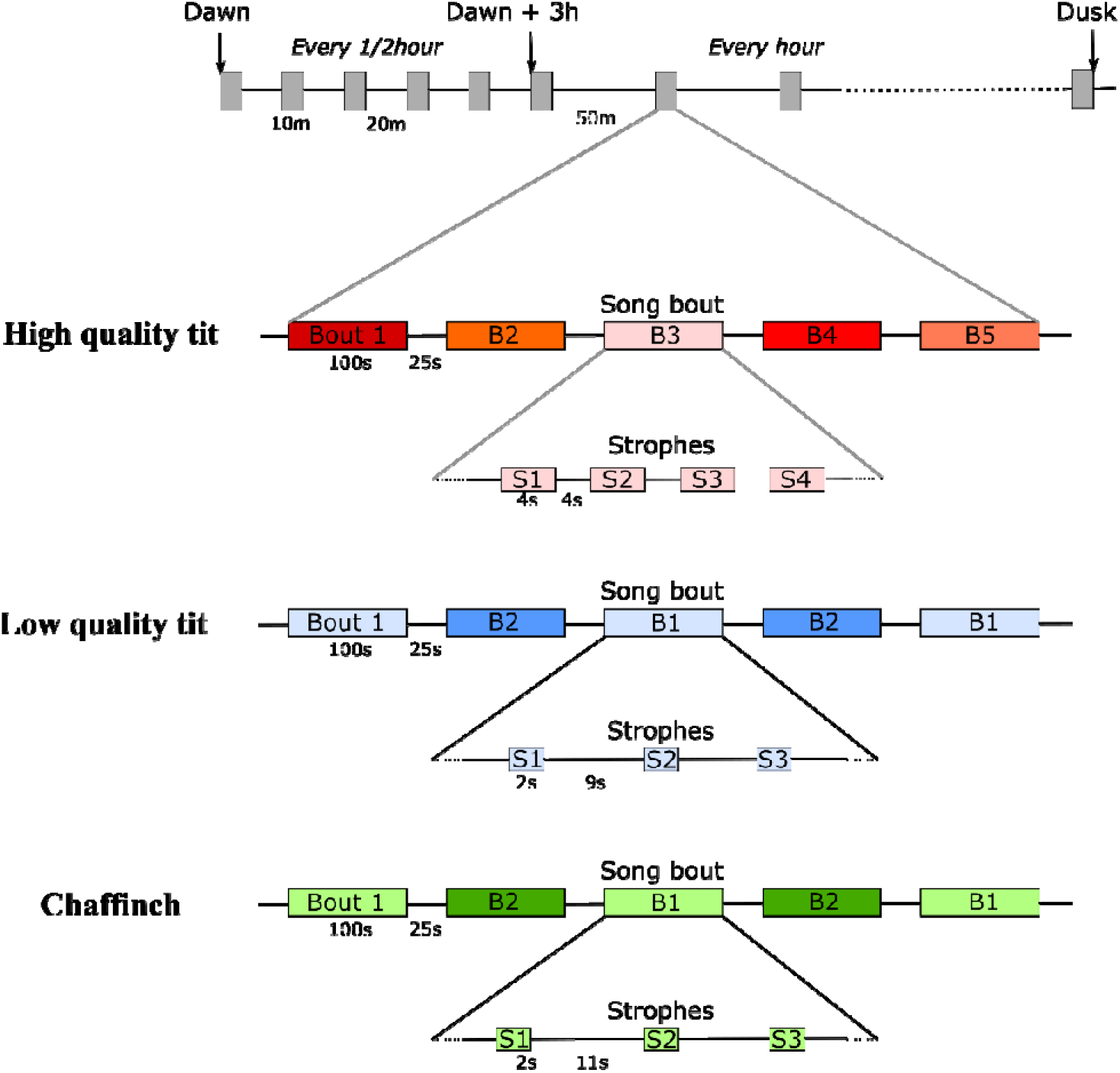
Structure of a song track (top), composed of a succession of 10 minute-long song periods. Song periods are composed of song bouts (B1, B2…), which are composed of strophes (S1, S2 …). All strophes in the same song bout were composed of the same type of syllable. Depending on the great tit natural song used for building the song track, the syllables within a strophe slightly varied in rhythm, amplitude, and, before transformation, in length. To standardize strophe length, we duplicated or deleted syllables. (A) Playback tracks mimicking a good quality great tit song had a repertoire size of 5 song bout types, composed of 4 second-long strophes separated by 4 seconds of silence. (B) Playback tracks mimicking a low quality song had a repertoire size of 2 song bout types, composed of 2 second-long strophes separated by 9 seconds of silence. The order of song bouts within a song period – and of strophes within song bouts- were alternated between song periods and song bouts to avoid habituation. (C) Chaffinch song track followed the same temporal pattern than low quality tit treatment (but with one strophe every 11 seconds), as it better matches their natural singing rhythm. The two different chaffinch song bout types per individual B1 and B2 were composed of a fixed syllable structure that could vary between individuals but was quite conserved within individuals.

